# Shared-Encoder Residual Task-Bridging Network for Motor Execution to Motor Imagery EEG Transfer in Brain-Computer Interfaces

**DOI:** 10.64898/2026.11.17.767217

**Authors:** Pengpai Wang, Yuxuan Zhu, Tiantian Xie, Wenlong Shen, Yueying Zhou, Xiaorui Zhang

## Abstract

Motor-imagery brain-computer interfaces (MI-BCIs) hold promising application prospects in the fields of limb motor rehabilitation and human-computer interaction. However, the reliability of MI calibration data is limited by the difficulty of verifying participants’ engagement during motor imagery whereas motor execution (ME) provides a more readily supervised source of motor-related information.To investigate the potential of ME-derived knowledge for assisting MI decoding, we propose a Shared-Encoder Residual Task-Bridge Network (SE-RTBNet) to achieve accurate intra-subject cross-task knowledge transfer from ME to MI. The proposed frame-work consists of two major components: a shared encoder and a residual task-bridge module. The shared encoder extracts stable general motor representations from multi-band EEG signals and explores the shared feature patterns underlying ME and MI tasks. The residual task-bridge module is equipped with a performance-aware routing mechanism. It adaptively compensates for task-specific discrepancies between the two tasks, enabling precise cross-task feature alignment and knowledge transfer. To validate the effectiveness of the proposed model, experiments are conducted on three public datasets. The proposed method achieves average classification accuracies of 69.39%, 74.61%, and 98.94%, respectively, and outperforms existing transfer-learning algorithms in overall performance. Our model provides an effective strategy for reducing dependence on extensive MI calibration data through intra-subject cross-task knowledge transfer.The source code of the proposed SE-RTBNet is publicly available at https://github.com/Daisyzyx2002/SE-RTBNet.

## I. Introduction

BRAIN-computer interface (BCI) establishes a direct neural communication pathway between the human brain and external devices, offering a reliable technical solution for clinical motor rehabilitation and human-computer interaction [1], [2]. Among various non-invasive BCI paradigms, electroencephalography (EEG)-based motor imagery (MI)-BCI has been extensively investigated owing to its portability, low cost, and non-invasive nature [3]–[7]. This technology decodes human motor intentions from modulations of sensorimotor cortical rhythms elicited by voluntary motor imagery [8], [9], holding great application potential for motor function rehabilitation of stroke survivors and assistive interaction for individuals with motor disorders. Nevertheless, the reliability of MI-BCI systems remains challenging to ensure in practical applications, partly because participants’ actual engagement during voluntary motor imagery is difficult to objectively verify. Insufficient or inconsistent engagement may result in unreliable MI calibration data, thereby affecting feature learning and decoding performance. In contrast, motor execution (ME) involves observable physical movements, providing a more readily supervised source of motor-related information. Neurophysiological evidence confirms that ME and MI tasks share highly overlapping activation patterns and spectral modulation characteristics within the sensorimotor cortex [10]–[12]. This indicates that ME data contain complementary knowledge of general motor representations, which can potentially support MI feature learning and decoding. Therefore, leveraging intra-subject ME data to assist MI decoding through cross-task knowledge transfer offers a promising strategy for reducing dependence on extensive labeled MI calibration data.

To mitigate the heavy data dependence and poor individual adaptability of conventional MI decoding pipelines, transfer learning has been widely adopted in BCI to reduce calibration overhead and improve model generalization [13], [14]. Existing transfer learning studies for BCI mostly focus on cross-subject and cross-session domain adaptation, with mainstream approaches including Riemannian manifold alignment [15]– [18], Euclidean space calibration [19], [20], optimal transport-based distribution adaptation [21]–[23], and deep adversarial domain adaptation [24]–[26]. Traditional inter-subject MI-to-MI transfer is constrained by inter-individual EEG variability and suffers limited adaptability, additionally, it overlooks the physiological mechanisms and spectral characteristics distinguishing ME and MI within the same subject. By contrast, intra-subject ME-to-MI transfer enjoys stronger consistency in homologous motor representations and more controllable residual discrepancies, enabling more stable cross-task transfer. Existing methods fail to exploit this advantage and tend to suffer from task-ambiguous feature entanglement and sample-level negative transfer. Furthermore, most existing studies neglect the inherent cross-task complementarity between ME and MI at the intra-subject level.

Compared with within-task motor imagery transfer, which has a well-established research foundation, studies focusing on within-subject cross-task transfer from motor execution to motor imagery remain relatively limited. Only a few existing works have carried out preliminary explorations. For instance, Lee et al. previously proposed a Siamese relational network and verified that ME could facilitate upper-limb MI decoding [27]. However, their work only exploited task similarity without modeling task-specific discrepancies. Subsequent channel-wise VAE-CNN frameworks validated that ME representations reconstructed alongside a small number of MI samples could reduce calibration costs, but they were confined to global feature adaptation and incapable of disentangling task-invariant information and task-specific residuals [28]. Miao et al. presented an interpretable cross-task transfer method that performed decoding via ME pre-training followed by limited MI fine-tuning [29]. Nevertheless, it relied on global alignment and lacked mechanisms to identify informative ME representations for transfer, limiting its ability to support MI decoding under limited calibration data conditions. Gwon et al. investigated cross-task transfer among ME, MI, and motor observation, demonstrating the potential of ME as the source task [30]. Still, their approach only adopted model transfer and few-shot fine-tuning without separating shared and task-specific representations, yielding only modest performance improvements under zero-shot settings. Pérez-Velasco et al. explored whether ME-pre-trained models could directly decode MI without MI fine-tuning. Interpretability analyses demonstrated that ME and MI EEG features are not equivalent, and ideal transfer cannot be achieved merely through global feature sharing [31]. Wang et al. developed a domain-class hierarchical adaptation network for intra-subject ME-to-MI EEG cross-task transfer decoding [32], but it only performed domain-level reweighting and lacked trial-wise assessment of ME sample transferability, failing to suppress negative transfer at the single-trial level. In summary, although existing ME-to-MI cross-task transfer methods have validated the feasibility of leveraging motor execution data to aid motor imagery decoding, critical limitations persist. Most methods depend on global feature alignment, representation reconstruction, or pre-training plus fine-tuning. They neither explicitly model task-specific residual differences between ME and MI nor account for the transfer heterogeneity of ME samples, and thus cannot adaptively identify transferable features.

To address the above limitations, this paper proposes a Shared Encoder Residual Task-Bridging Network (SE-RTBNet) for adaptive intra-subject cross-task transfer learning of EEG signals from motor execution to motor imagery. The framework mainly consists of a shared encoder and a residual task-bridging module with performance-aware routing. The shared encoder conducts deep feature extraction on multi-band EEG signals to capture stable, generalizable sensorimotor representations shared by ME and MI, preserving shared motor-related information across the two tasks. On this basis, the residual task-bridging module discards the rigid global feature alignment constraints widely adopted in conventional methods. It employs learnable residual mappings to explicitly characterize task-specific discrepancies between ME and MI, separating task-invariant shared features from task-specific differentiated features. Meanwhile, the embedded performance-aware routing mechanism adaptively fuses multiple task-bridging branches to accommodate transfer heterogeneity across distinct motor categories and precisely compensate for feature biases between tasks. This breaks the limitations of conventional global adaptation strategies, enabling explicit characterization of inherent task-specific differences between ME and MI while adaptively screening transferable features according to the heterogeneity of ME samples.

The main contributions of this paper are summarized as follows:

1. We propose a shared representation learning strategy that integrates multi-band sensorimotor encoding, distribution alignment, supervised contrastive learning, and spatio-temporal regularization to improve the robust discriminative capability of ME-to-MI EEG features.
2. We develop a residual task bridging framework and multi-expert performance-aware routing mechanism for ME-to-MI transfer. By formulating cross-task feature shift as learnable residual mappings, the framework circumvents the limitations of traditional global feature alignment and adapts to transfer heterogeneity via adaptive fusion of multiple task-bridging branches.
3. We conduct extensive comparative and ablation experiments on public EEG datasets. Comparisons between SE=RTBNet and recent state-of-the-art methods validate the efficacy of our model and facilitate the practical deployment of lightweight, highly adaptable MI-BCI systems.

## II. Materials and methods

### A. Datasets and Preprocessing

To validate the effectiveness and cross-dataset generalization of the proposed SE-RTBNet for ME-to-MI transfer learning, this study employs three public EEG datasets containing paired ME and MI paradigms. The dataset details and corresponding preprocessing pipelines are presented below.

#### 1) PhysioNet EEG Motor Movement/Imagery Dataset

This study uses the public PhysioNet EEG Motor Movement/Imagery Dataset (Physionet Dataset) as the benchmark for cross-task transfer learning [33]. Recorded via the BCI2000 system from 109 healthy subjects, the dataset adopts 64 electrodes following the international 10-10 layout at an original sampling rate of 160 Hz. Designed in blocks, it comprises real movement, motor imagery and resting-state paradigms over multiple independent runs. We select left- and right-hand tasks for binary cross-task validation: ME samples come from runs 3, 7 and 11, whereas MI samples are taken from runs 4, 8 and 12. Both paradigms have a 4 s single-trial duration with accurate event markers.

Given Subjects 88, 92 and 100 were originally sampled at 128 Hz and others at 160 Hz, all recordings were resampled to 160 Hz with anti-aliasing interpolation and standardized for channel montage. Continuous EEG was zero-phase high-pass filtered at 1 Hz, notch-filtered at 60 Hz for baseline drift and line noise removal, then re-referenced to CAR. After epoching by event markers and baseline correction over [−1.0, 0] s pre-cue, the stable 0.5-3.5 s post-cue segment was retained for modeling. We selected 21 sensorimotor channels (FC1-FC6, C1-C6, Cz, CP1-CP6, P3, P4) to reduce irrelevant noise.

#### 2) Multi-paradigm Upper Limb Rehabilitation EEG Dataset

This study adopts the multi-paradigm upper-limb rehabilitation EEG dataset (MULR Dataset) proposed by Chang et al. [34]. It contains recordings from 28 right-handed healthy subjects (14 males, 14 females, aged 20-33). EEG was sampled at 1000 Hz via 59 scalp electrodes arranged under the international 10-20 system; auxiliary EOG and ECG were recorded for artifact rejection. Among its six rehabilitation paradigms, we select only pure motor execution and motor imagery for cross-task transfer. Subjects performed or imagined left-hand, right-hand and bimanual grasp-and-release movements guided by visual cues. Each trial comprises a 4 s task period and a 2 s rest interval, and only left- and right-hand trials are retained for analysis.

Continuous EEG from the four runs of each paradigm is concatenated and mapped to the standard 10-05 montage. A zero-phase 0.1-40 Hz bandpass filter removes baseline drift and high-frequency noise, followed by downsampling to 160 Hz for uniform preprocessing. Epochs are segmented by event markers, baseline-corrected over [™0.5, 0] s pre-cue, and cropped to the 0.5-3.5 s post-cue steady task window. The same 21 sensorimotor channels as Dataset 1 are preserved to isolate motor cortical activity. Subjects 207 and 218 are excluded owing to recording anomalies.

#### 3) Simultaneous EEG-fMRI Motor Execution/Imagery Dataset

This study adopts the simultaneous EEG-fMRI motor-task dataset (SEMI Dataset) from Bondi et al. [12]. Of the initially recruited 37 healthy subjects who completed 3-5 MI training sessions for one-dimensional cursor control, 17 passed screening (valid accuracy 70% and full participation) for the simultaneous EEG-fMRI recording. EEG was captured via 63 electrodes under the international 10-20 system at 5000 Hz. The block-designed experiment alternated task and rest periods with ME and MI paradigms. We retained only left- and right-hand fist-clenching tasks cued at 2 Hz. For each action, ME blocks preceded MI blocks, while the global block order was randomized and balanced across subjects.

We used the authors’ pre-cleaned EEG, downsampled from 250 Hz to 160 Hz. Built-in stimulus markers separated ME and MI trials, and labels were standardized for binary left/right-hand classification. We extracted 0-3 s post-onset epochs and retained the same 21 sensorimotor channels for consistent spatial representation.

### B. Overall of Method

The proposed SE-RTBNet first performs multi-band decomposition and feature alignment. It then extracts shared feature patterns of ME and MI via a shared encoder. Afterward, the residual task-bridging module adaptively compensates for task-specific discrepancies between ME and MI. Finally, the model matches features from five sliding temporal windows of test samples against bridged prototypes and obtains classification results with weighted test-time augmentation. The overall architecture of the proposed SE-RTBNet is illustrated in Fig. 1.

**Fig. 1.**
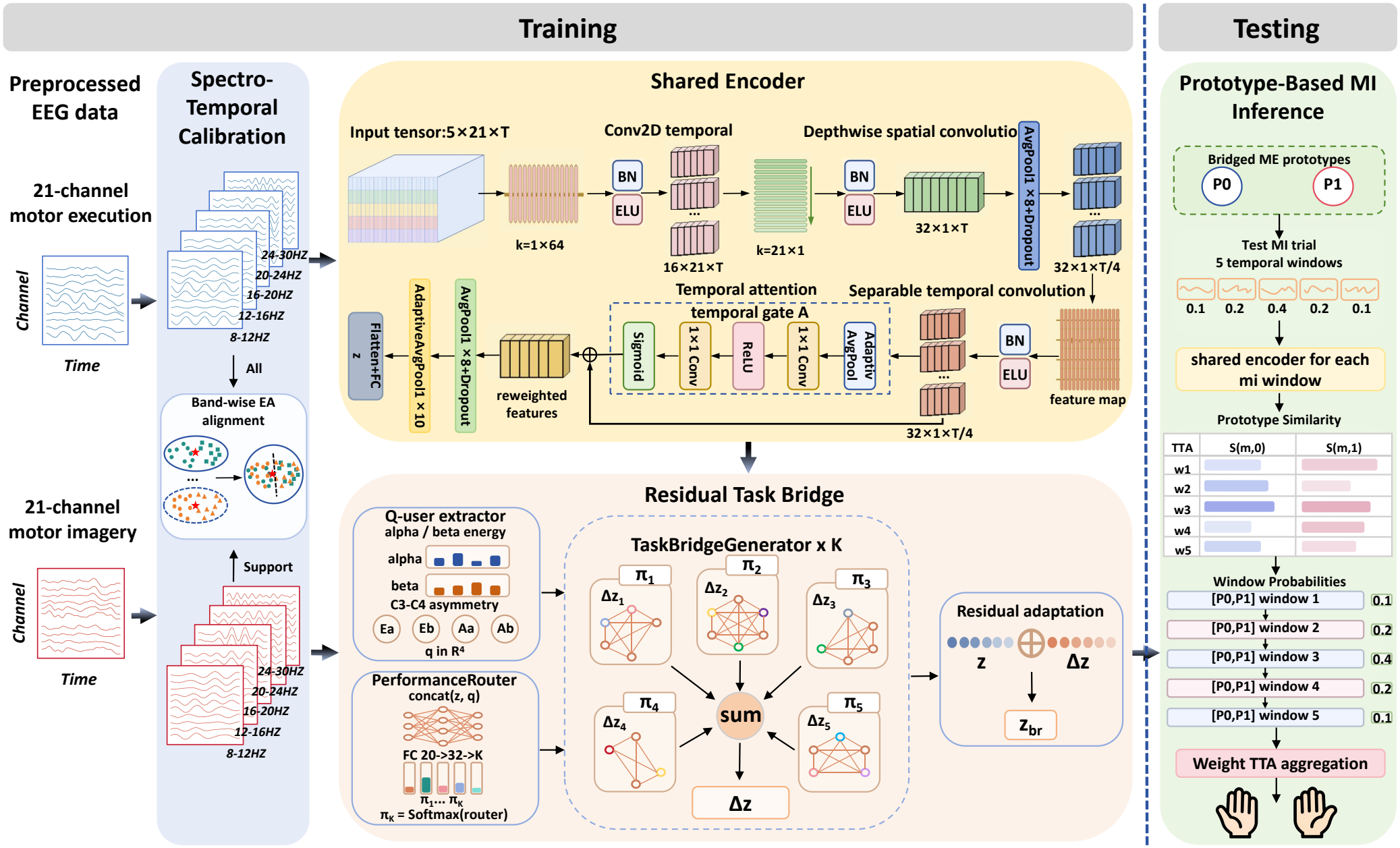
Architecture of the proposed SE-RTBNet.During the training phase, SE-RTBNet first applies multi-band decomposition and Euclidean alignment to calibrate the distributions of ME and MI signals. A shared encoder is then employed to extract task-invariant representations shared between ME and MI, followed by a residual task-bridge module that adaptively compensates for task-specific discrepancies. During the testing phase, each test trial is partitioned into five sliding temporal windows to extract latent representations, which are evaluated against the transfer-calibrated class prototypes via similarity matching. Finally, a weighted test-time augmentation (TTA) strategy is incorporated to aggregate multi-window prediction probabilities, yielding the binary classification decision.

#### 1) Spectro-Temporal Calibration

To evaluate the proposed framework, a stratified five-fold cross-validation scheme is conducted on the MI dataset. In each fold, one MI subset is held out as the test set *D*_te_, while the remaining MI trials are partitioned into a support subset 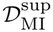 and a validation subset 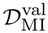. All ME trials _ME_ are leveraged as source-task training data across all folds.

In the spectro-temporal calibration stage, preprocessed EEG trials are first decomposed via a filter bank covering *B* sub-bands. Each trial is represented as a tensor 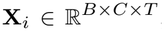, where *B, C*, and *T* denote the number of frequency bands, recording channels, and time samples, respectively. Multi-band Euclidean Alignment (EA) is applied independently to the spatial slice **X**_*i,b*_ ∈ ℝ^*C*×*T*^ of each frequency band *b* ∈ {1, … , *B*} to mitigate cross-task covariance distribution shifts:

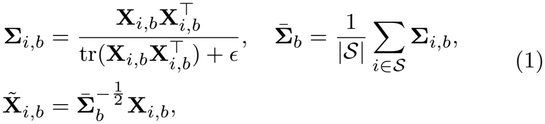

where *ϵ >* 0 is a regularization constant preventing numerical instability. The alignment reference matrix 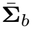 for the ME domain is estimated across all ME trials (*S*= *D*_ME_), whereas that for the MI domain is estimated strictly from the current MI support subset 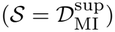 to prevent data leakage.

From each aligned sample 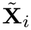, a four-dimensional neurophysiological descriptor **q**_*i*_ is extracted to quantify spectral power and sensorimotor rhythm lateralization:

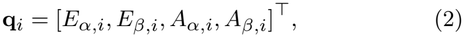

where 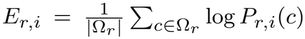 represents the log-band power averaged over the central sensorimotor channels Ω_*r*_ in rhythm *r* ∈ {*α, β*} . The hemispheric lateralization index *A*_*r,I*_ is formulated as:

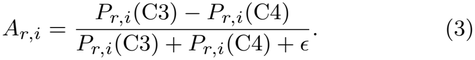

#### 2) Shared Encoder Model

The shared encoder *f*_*θ*_ takes the multi-band tensor 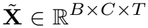 as input and maps it into a low-dimensional latent space 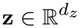 . Rather than employing separate parallel branches, the *B* sub-bands are treated as input feature planes for 2D spatio-temporal convolutions, enabling early cross-band feature fusion with a minimal parameter footprint.

The feature extractor adopts a temporal-spatial-temporal cascade:

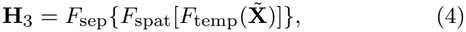

where *F*_temp_ denotes a 2D temporal convolution with a kernel size of 1 × *K*_*t*_ (capturing phase dynamics within sub-bands), *F*_spat_ is a depthwise spatial convolution with a kernel size of *C* 1 parameterized by spatial filter weights 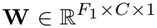, and *F*_sep_ is a separable temporal convolution with a kernel size of 1 × *K*_*s*_ for fine-grained temporal abstraction. Each block incorporates Batch Normalization, Exponential Linear Unit (ELU) activations, Average Pooling, and Dropout.

To adaptively highlight task-relevant ERD/ERS periods, a temporal squeeze-and-excitation attention module is appended after **H**_3_:

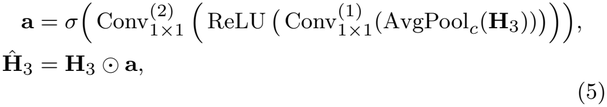

where AvgPool_*c*_(·) squeezes the spatial/channel dimensions via average pooling, Conv_1×1_ layers act as temporal excitation bottlenecks, and *σ*(·) denotes the Sigmoid function producing the temporal gating vector 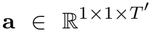 . The calibrated feature tensor **Ĥ**_3_ is flattened and projected via a linear layer:

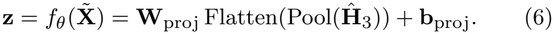

The shared encoder and auxiliary classifier are optimized end-to-end via a multi-task joint objective, where individual losses are dynamically balanced via homoscedastic uncertainty with learnable log-variance parameters *s*_*k*_ :

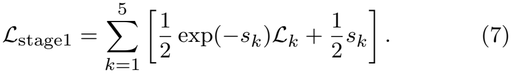

The individual loss components are formulated as follows:

a. ME Mixup Classification Loss (ℒ_1_): To regularize source-task representations and mitigate overfitting to subject-specific artifacts, Mixup data augmentation is applied to the aligned ME trials:

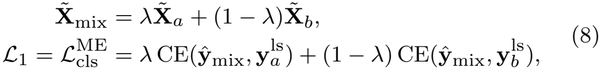

where 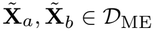 are randomly paired source trials within the training batch, *λ* ∼ Beta(*α, α*) is the interpolation ratio, and 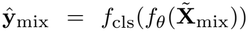 represents the predicted class probabilities. To further prevent overconfident predictions, one-hot labels **y** are converted into smoothed targets **y**^ls^ = (1 *ϵ*_ls_)**y**+*ϵ*_ls_*/N*_*c*_, where *ϵ*_ls_ is the label smoothing factor, *N*_*c*_ = 2 is the number of classes, and CE(**p, q**) = *q*_*c*_ log *p*_*c*_ denotes the cross-entropy function.
b. MI Support Classification Loss (ℒ_2_): To retain class discriminability on target representations, standard cross-entropy is computed over the few-shot MI support subset:

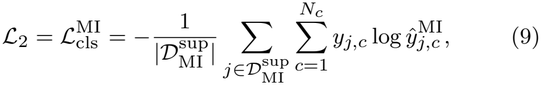

where *y*_*j,c*_ ∈ {0, 1} is the ground-truth class indicator, and 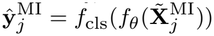.
c. Multi-Kernel MMD Loss (ℒ_3_): To reduce cross-task domain discrepancy in the latent space, multi-kernel maximum mean discrepancy (MK-MMD) aligns marginal distributions between the source ME domain and the target MI support subset:

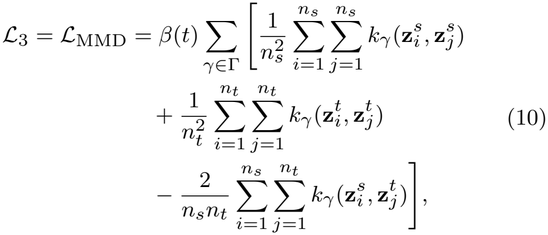

where 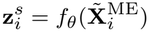 and 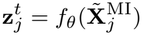 denote latent features from source and target batches of sizes *n*_*s*_ and *n*_*t*_, respectively. The Gaussian kernel is 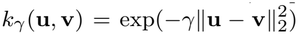, with Γ denoting the set of bandwidths. The indicator *β*(*t*) = I(*t > E*_warm_) denotes a warmup schedule at epoch *t* to prevent premature representation distortion.
d. Supervised Contrastive Loss (ℒ_4_): To enforce cross-task semantic alignment, this loss pulls same-class ME and MI pairs together while pushing discordant classes apart:

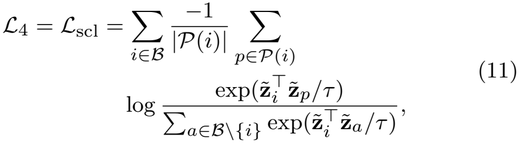

where 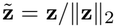 denotes the ℓ_2_-normalized feature, *B* is the combined ME–MI mini-batch, *P*(*i*) = {*p* ∈ *B\* {*i*} : *y*_*p*_ = *y*_*i*_}indexes cross-task positive samples, and *τ >* 0 is the temperature parameter.
e. Neurophysiological Regularization (ℒ_5_): Domain priors regarding cortical spatial topology and temporal dynamics are enforced via:

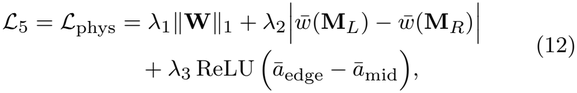

where 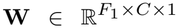 denotes the spatial filter weights, and 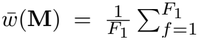 mean(|**W**_*f*_ | ⊙ **M**) computes the average weight activation over left/right motor cortex masks **M**_*L*_, **M**_*R*_ {0, 1} ^*C*×1^. For temporal attention 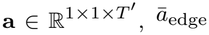 , *ā*_edge_ and *ā*_mid_ correspond to the mean activations over edge windows 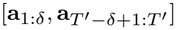 and the central execution window 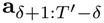 , penalizing spurious attention on non-task baseline intervals.

#### 3) Residual Task Bridge Model

Although the shared encoder captures task-invariant sensorimotor features, task-specific distribution discrepancies persist between ME and MI. To bridge this residual shift, we propose a residual task bridging network governed by a performance-guided routing mechanism, which maps source ME representations toward the MI target manifold in a residual additive manner.

Given the latent representation 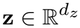 and the neurophysiological descriptor **q** ℝ^4^, the router predicts sample-adaptive gating coefficients across *K* specialized bridge experts:

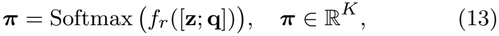

where *f*_*r*_ denotes the multilayer perceptron of the router, and [**z**; **q**] represents vector concatenation.

Each bridge expert 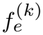 outputs an expert-specific residual correction vector:

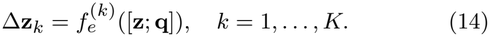

The aggregated residual vector is obtained through gate-weighted summation, yielding the bridged representation **z**_br_:

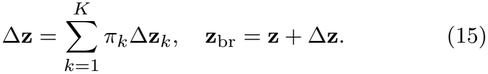

Once Stage 1 training is accomplished, the shared encoder *f*_*θ*_ is frozen. The bridge experts and router are optimized via a class-wise prototype alignment loss in Stage 2. First, the bridged ME class prototypes and target MI support prototypes are computed:

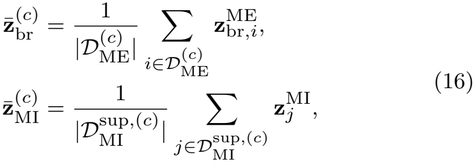

where 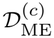 and 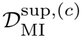 denote class-*c* samples from the source ME and target MI support sets, respectively.

The bridging loss minimizes the squared Euclidean distance between corresponding class prototypes, guiding the bridged ME representations toward the empirical MI distribution:

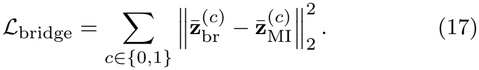

During test inference, task prototypes are derived from all available ME samples transformed by the trained bridging module:

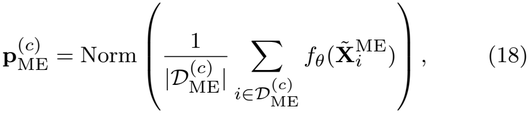

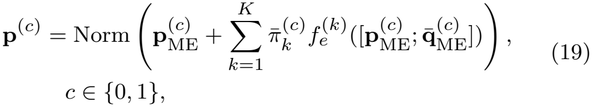

where 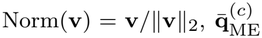 denotes the class-conditional average descriptor over source trials, and 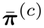 represents the empirical average routing weights for class *c*, yielding the bridged target prototypes **p**^(0)^ and **p**^(1)^.

For an unseen MI test trial 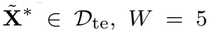 overlapping temporal windows {*w*_1_, … , *w*_*W*_} are extracted via test-time augmentation (TTA). For the *w*-th temporal window, the normalized latent representation 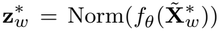 is extracted. The prototype similarity score *S*(*w, c*) with respect to class prototype **p**^(*c*)^ is computed as:

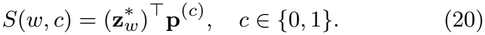

The window-level class probability vector **r**_*w*_ ℝ^2^ is then evaluated using scaled cosine logits:

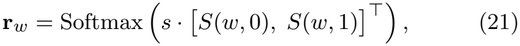

where *s* = 10 is an inverse temperature parameter scaling prediction confidence.

Finally, multi-window predictions are aggregated using a symmetric pyramidal weight vector ***ω*** = [*ω*_1_, … , *ω*_*W*_]^⊤^ = [0.1, 0.2, 0.4, 0.2, 0.1]^⊤^:

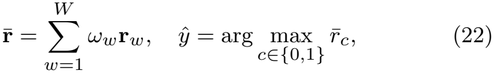

where 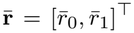 denotes the ensemble probability vector, and *ŷ* is the predicted class label.

## III. Results

Our proposed SE-RTBNet addresses ME-to-MI cross-task transfer decoding. Four experiments are designed to evaluate its performance. The model is compared with existing methods to validate its effectiveness. Within-task MI decoding under identical dataset conditions, subject partitions, and classifier configurations quantifies the performance gap between cross-task and within-task decoding. Subject-level accuracy distributions across three benchmark datasets characterize inter-subject variability and generalization capability. Hyperparameter tuning further examines the effects of key architectural parameters on overall performance.

### A. Cross-Task Transfer Performance

The cross-task transfer scenario from ME to MI serves as the primary evaluation benchmark of this study. For rigorous and objective comparisons across diverse methodological paradigms, ten representative baseline models are adopted: three classical spatial-filtering pipelines (CSP+SVM [35], CSP+KNN [35], FBCSP+SVM [36]), four mainstream deep-learning decoders (EEGNet [37], ShallowConvNet [38], DeepConvNet [38], EEGConformer [3]), and three BCI-oriented cross-task transfer-learning models (CT-adaptTL [29], EEGSym [39], CVNet [28]). Results on the three datasets are summarized in Table I.

**TABLE 1.** Cross-task transfer performance from motor execution to motor imagery across three benchmark datasets. Values are reported as mean ± interpatient standard deviation. Best results are highlighted in bold. Statistical significance relative to se-rtbnet is indicated as ^*^ (***p <* 0.05**), ^**^ (***p <* 0.01**), and ^***^ (***p <* 0.001**).

| Dataset | Model | Accuracy (%) | AUC | F1 Score | Recall |
|---|---|---|---|---|---|
| PhysioNet Dataset | CSP+SVM [35] | 63.69<sub>±21.21</sub><sup>***</sup> | 0.677<sub>±0.249</sub><sup>**</sup> | 0.621<sub>±0.248</sub><sup>***</sup> | 0.653<sub>±0.288</sub><sup>**</sup> |
|  | CSP+KNN [35] | 63.68<sub>±21.05</sub><sup>***</sup> | 0.675<sub>±0.247</sub><sup>***</sup> | 0.620<sub>±0.245</sub><sup>***</sup> | 0.645<sub>±0.280</sub><sup>**</sup> |
|  | FBCSP+SVM [36] | 63.51<sub>±20.39</sub><sup>***</sup> | 0.684<sub>±0.237</sub><sup>**</sup> | 0.622<sub>±0.230</sub><sup>***</sup> | 0.647<sub>±0.273</sub><sup>***</sup> |
|  | EEGNet [37] | 63.71<sub>±20.41</sub><sup>***</sup> | 0.674<sub>±0.244</sub><sup>***</sup> | 0.620<sub>±0.235</sub><sup>*</sup> | 0.641<sub>±0.271</sub><sup>***</sup> |
|  | ShallowConvNet [38] | 61.51<sub>±20.13</sub><sup>***</sup> | 0.652<sub>±0.244</sub><sup>***</sup> | 0.594<sub>±0.238</sub><sup>***</sup> | 0.614<sub>±0.274</sub><sup>***</sup> |
|  | EEGConformer [3] | 64.14<sub>±20.27</sub><sup>***</sup> | 0.683<sub>±0.241</sub><sup>**</sup> | 0.618<sub>±0.244</sub><sup>***</sup> | 0.636<sub>±0.279</sub><sup>***</sup> |
|  | DeepConvNet [38] | 63.43<sub>±19.66</sub><sup>***</sup> | 0.670<sub>±0.239</sub><sup>***</sup> | 0.610<sub>±0.240</sub><sup>**</sup> | 0.631<sub>±0.280</sub><sup>***</sup> |
|  | CT-adaptTL [29] | 61.97<sub>±14.09</sub><sup>***</sup> | 0.652<sub>±0.158</sub><sup>***</sup> | 0.593<sub>±0.156</sub><sup>***</sup> | 0.621<sub>±0.140</sub><sup>***</sup> |
|  | EEGSym [39] | 63.64<sub>±14.34</sub><sup>***</sup> | 0.680<sub>±0.176</sub><sup>***</sup> | 0.612<sub>±0.175</sub><sup>***</sup> | 0.636<sub>±0.199</sub><sup>***</sup> |
|  | CVNet [28] | 65.92<sub>±8.73</sub><sup>***</sup> | 0.560<sub>±0.158</sub><sup>***</sup> | 0.637<sub>±0.135</sub><sup>**</sup> | **0.713<sub>±0.170</sub>** |
|  | **SE-RTBNet** (**Ours**) | **69.39<sub>±9.41</sub>** | **0.715<sub>±0.136</sub>** | **0.672<sub>±0.107</sub>** | 0.694<sub>±0.096</sub> |
| MULR Dataset | CSP+SVM [35] | 70.65<sub>±14.34</sub><sup>*</sup> | 0.750<sub>±0.167</sub><sup>*</sup> | 0.695<sub>±0.168</sub> | 0.704<sub>±0.192</sub> |
|  | CSP+KNN [35] | 68.20<sub>±14.52</sub><sup>***</sup> | 0.723<sub>±0.157</sub><sup>***</sup> | 0.679<sub>±0.149</sub><sup>**</sup> | 0.687<sub>±0.154</sub><sup>*</sup> |
|  | FBCSP+SVM [36] | 70.91<sub>±15.38</sub><sup>*</sup> | 0.752<sub>±0.174</sub><sup>*</sup> | 0.702<sub>±0.162</sub><sup>*</sup> | 0.712<sub>±0.161</sub> |
|  | EEGNet [37] | 60.89<sub>±10.68</sub><sup>***</sup> | 0.640<sub>±0.131</sub><sup>***</sup> | 0.591<sub>±0.134</sub><sup>**</sup> | 0.610<sub>±0.152</sub><sup>**</sup> |
|  | ShallowConvNet [38] | 61.04<sub>±10.92</sub><sup>***</sup> | 0.656<sub>±0.139</sub><sup>***</sup> | 0.594<sub>±0.154</sub><sup>***</sup> | 0.643<sub>±0.181</sub><sup>**</sup> |
|  | EEGConformer [3] | 69.32<sub>±13.93</sub><sup>*</sup> | 0.764<sub>±0.149</sub> | 0.693<sub>±0.142</sub><sup>*</sup> | 0.726<sub>±0.136</sub> |
|  | DeepConvNet [38] | 61.16<sub>±10.60</sub><sup>***</sup> | 0.638<sub>±0.130</sub><sup>***</sup> | 0.593<sub>±0.118</sub><sup>***</sup> | 0.618<sub>±0.124</sub><sup>***</sup> |
|  | CT-adaptTL [29] | 68.33<sub>±11.93</sub><sup>***</sup> | 0.750<sub>±0.134</sub><sup>*</sup> | 0.678<sub>±0.121</sub><sup>***</sup> | 0.683<sub>±0.119</sub><sup>***</sup> |
|  | EEGSym [39] | 71.19<sub>±14.22</sub><sup>**</sup> | 0.772<sub>±0.163</sub> | 0.688<sub>±0.163</sub><sup>**</sup> | 0.692<sub>±0.183</sub><sup>*</sup> |
|  | CVNet [28] | 73.72<sub>±10.83</sub> | 0.721<sub>±0.145</sub><sup>*</sup> | 0.730<sub>±0.118</sub> | **0.758<sub>±0.126</sub>** |
|  | **SE-RTBNet** (**Ours**) | **74.61<sub>±13.87</sub>** | **0.789<sub>±0.152</sub>** | **0.740<sub>±0.144</sub>** | 0.746<sub>±0.139</sub> |
| SEMI Dataset | CSP+SVM [35] | 73.19<sub>±6.67</sub><sup>***</sup> | 0.809<sub>±0.071</sub><sup>***</sup> | 0.730<sub>±0.074</sub><sup>***</sup> | 0.732<sub>±0.096</sub><sup>***</sup> |
|  | CSP+KNN [35] | 85.47<sub>±3.87</sub><sup>***</sup> | 0.936<sub>±0.027</sub><sup>***</sup> | 0.855<sub>±0.039</sub><sup>***</sup> | 0.858<sub>±0.045</sub><sup>***</sup> |
|  | FBCSP+SVM [36] | 81.15<sub>±6.32</sub><sup>***</sup> | 0.887<sub>±0.060</sub><sup>***</sup> | 0.814<sub>±0.064</sub><sup>***</sup> | 0.826<sub>±0.072</sub><sup>***</sup> |
|  | EEGNet [37] | 82.49<sub>±7.96</sub><sup>***</sup> | 0.904<sub>±0.069</sub><sup>***</sup> | 0.829<sub>±0.077</sub><sup>***</sup> | 0.847<sub>±0.077</sub><sup>***</sup> |
|  | ShallowConvNet [38] | 94.47<sub>±2.17</sub><sup>***</sup> | 0.986<sub>±0.009</sub><sup>***</sup> | 0.945<sub>±0.022</sub><sup>***</sup> | 0.946<sub>±0.022</sub><sup>***</sup> |
|  | EEGConformer [3] | 85.00<sub>±5.84</sub><sup>***</sup> | 0.925<sub>±0.047</sub><sup>***</sup> | 0.850<sub>±0.059</sub><sup>***</sup> | 0.850<sub>±0.066</sub><sup>***</sup> |
|  | DeepConvNet [38] | 94.26<sub>±2.28</sub><sup>***</sup> | 0.988<sub>±0.008</sub><sup>***</sup> | 0.943<sub>±0.022</sub><sup>***</sup> | 0.949<sub>±0.024</sub><sup>***</sup> |
|  | CT-adaptTL [29] | 86.60<sub>±5.70</sub><sup>***</sup> | 0.930<sub>±0.045</sub><sup>***</sup> | 0.866<sub>±0.057</sub><sup>***</sup> | 0.866<sub>±0.057</sub><sup>***</sup> |
|  | EEGSym [39] | 93.49<sub>±2.95</sub><sup>***</sup> | 0.979<sub>±0.017</sub><sup>***</sup> | 0.935<sub>±0.030</sub><sup>***</sup> | 0.934<sub>±0.031</sub><sup>***</sup> |
|  | CVNet [28] | 93.96<sub>±3.78</sub><sup>***</sup> | 0.984<sub>±0.016</sub><sup>***</sup> | 0.940<sub>±0.036</sub><sup>***</sup> | 0.947<sub>±0.047</sub><sup>**</sup> |
|  | **SE-RTBNet** (**Ours**) | **98.94<sub>±0.74</sub>** | **0.999<sub>±0.001</sub>** | **0.989<sub>±0.007</sub>** | **0.989<sub>±0.007</sub>** |

On the PhysioNet dataset, conventional deep networks and spatial-filtering pipelines suffer substantial performance degradation, with classification accuracies ranging merely from 61.51% to 64.14%. Although the recently proposed transfer-learning baseline CVNet achieves a decent accuracy of 65.92%, SE-RTBNet yields an accuracy of 69.39%, outperforming CVNet and EEGConformer by 3.47 and 5.25 percentage points, respectively. Furthermore, SE-RTBNet greatly reduces the subject-level standard deviation to 9.41%, in contrast to approximately 20% for most conventional decoders. Even though CVNet attains a slightly lower standard deviation of 8.73%, its low AUC of 0.560 indicates that such low variance stems from conservative decision bias induced by forced domain alignment. By contrast, SE-RTBNet substantially suppresses variance while delivering genuine robustness with high accuracy and discriminative potential.

On the MULR dataset, the performance differences among competing methods become relatively narrow, indicating a more challenging cross-task transfer scenario. Although CVNet achieves competitive performance, with an accuracy of 73.72% and an F1-score of 0.730, SE-RTBNet further improves the overall decoding performance, attaining the highest accuracy of 74.61% and F1-score of 0.740. More notably, SE-RTBNet achieves an AUC of 0.789, substantially outperforming CVNet (0.721), with the improvement reaching statistical significance (*p <* 0.05). This pronounced AUC improvement suggests that SE-RTBNet provides more effective class discrimination beyond the final classification threshold. Overall, these results demonstrate that SE-RTBNet maintains competitive recall while achieving superior accuracy, discriminative capability, and F1-score, highlighting its effectiveness in cross-task knowledge transfer on the MULR dataset.

On the SEMI dataset, ShallowConvNet and DeepConvNet achieve accuracies of 94.47% and 94.26%, respectively, while CVNet records 93.96%. Remarkably, SE-RTBNet attains a near-saturated accuracy of 98.94% with a minimal subject-level standard deviation of 0.74%. Paired *t*-tests further corroborate that SE-RTBNet significantly outperforms all competing baselines on this dataset.

### B. Within-Task MI Decoding Performance

MI-to-MI within-task evaluations serve as the reference benchmark for quantifying the performance gap between ME-to-MI cross-task transfer and within-task MI decoding. Identical datasets, subject partitions and classifier configurations are adopted throughout these assessments. All baseline models are trained and tested on MI data, and corresponding experimental results are summarized in Table II.

**TABLE 2.** Classification performance of mi-trained and mi-tested models across three datasets, with our me-trained se-rtbnet included as a cross-task reference. Values are reported as mean ± inter-subject standard deviation. Best results are highlighted in bold.

| Dataset | Model | Accuracy (%) | AUC | F1 Score | Recall |
|---|---|---|---|---|---|
| PhysioNet Dataset | CSP+SVM [35] | 62.21<sub>±17.44</sub> | 0.647<sub>±0.212</sub> | 0.589<sub>±0.202</sub> | 0.600<sub>±0.197</sub> |
|  | CSP+KNN [35] | 62.13<sub>±17.49</sub> | 0.640<sub>±0.210</sub> | 0.592<sub>±0.198</sub> | 0.602<sub>±0.193</sub> |
|  | FBCSP+SVM [36] | 61.27<sub>±16.47</sub> | 0.644<sub>±0.200</sub> | 0.577<sub>±0.192</sub> | 0.587<sub>±0.195</sub> |
|  | EEGNet [37] | 71.41<sub>±9.18</sub> | 0.637<sub>±0.140</sub> | 0.654<sub>±0.156</sub> | 0.663<sub>±0.187</sub> |
|  | ShallowConvNet [38] | 71.88<sub>±11.23</sub> | 0.646<sub>±0.171</sub> | 0.667<sub>±0.164</sub> | 0.694<sub>±0.175</sub> |
|  | EEGConformer [3] | **79.20<sub>±10.20</sub>** | **0.723<sub>±0.152</sub>** | **0.748<sub>±0.153</sub>** | **0.762<sub>±0.168</sub>** |
|  | DeepConvNet [38] | 72.76<sub>±7.88</sub> | 0.650<sub>±0.123</sub> | 0.679<sub>±0.132</sub> | 0.693<sub>±0.154</sub> |
|  | SE-RTBNet(Ours) | 69.39<sub>±9.41</sub> | 0.715<sub>±0.136</sub> | 0.672<sub>±0.107</sub> | 0.694<sub>±0.096</sub> |
| MULR Dataset | CSP+SVM [35] | 67.87<sub>±15.90</sub> | 0.728<sub>±0.181</sub> | 0.672<sub>±0.168</sub> | 0.677<sub>±0.171</sub> |
|  | CSP+KNN [35] | 68.17<sub>±15.47</sub> | 0.715<sub>±0.168</sub> | 0.680<sub>±0.158</sub> | 0.693<sub>±0.159</sub> |
|  | FBCSP+SVM [36] | 70.91<sub>±17.02</sub> | 0.749<sub>±0.185</sub> | 0.709<sub>±0.169</sub> | 0.717<sub>±0.161</sub> |
|  | EEGNet [37] | 73.69<sub>±10.25</sub> | 0.708<sub>±0.137</sub> | 0.719<sub>±0.123</sub> | 0.735<sub>±0.124</sub> |
|  | ShallowConvNet [38] | 78.04<sub>±11.95</sub> | 0.770<sub>±0.148</sub> | 0.769<sub>±0.133</sub> | 0.796<sub>±0.129</sub> |
|  | EEGConformer [3] | **79.91<sub>±10.39</sub>** | 0.761<sub>±0.140</sub> | **794<sub>±0.113</sub>** | **0.807<sub>±0.128</sub>** |
|  | DeepConvNet [38] | 70.85<sub>±7.75</sub> | 0.669<sub>±0.099</sub> | 0.702<sub>±0.095</sub> | 0.725<sub>±0.125</sub> |
|  | SE-RTBNet (Ours) | 74.61<sub>±13.87</sub> | **0.789<sub>±0.152</sub>** | 0.740<sub>±0.144</sub> | 0.746<sub>±0.139</sub> |
| SEMI Dataset | CSP+SVM [35] | 84.00<sub>±5.32</sub> | 0.908<sub>±0.047</sub> | 0.840<sub>±0.055</sub> | 0.841<sub>±0.061</sub> |
|  | CSP+KNN [35] | 91.36<sub>±2.98</sub> | 0.971<sub>±0.015</sub> | 0.914<sub>±0.030</sub> | 0.916<sub>±0.037</sub> |
|  | FBCSP+SVM [36] | 89.86<sub>±5.51</sub> | 0.953<sub>±0.043</sub> | 0.898<sub>±0.055</sub> | 0.899<sub>±0.057</sub> |
|  | EEGNet [37] | 82.70<sub>±8.89</sub> | 0.888<sub>±0.083</sub> | 0.826<sub>±0.090</sub> | 0.833<sub>±0.092</sub> |
|  | ShallowConvNet [38] | 94.76<sub>±1.68</sub> | 0.987<sub>±0.007</sub> | 0.947<sub>±0.017</sub> | 0.946<sub>±0.023</sub> |
|  | EEGConformer [3] | 95.80<sub>±1.53</sub> | 0.993<sub>±0.005</sub> | 0.958<sub>±0.015</sub> | 0.961<sub>±0.019</sub> |
|  | DeepConvNet [38] | 93.96<sub>±2.57</sub> | 0.985<sub>±0.011</sub> | 0.940<sub>±0.026</sub> | 0.941<sub>±0.032</sub> |
|  | SE-RTBNet (Ours) | **98.94<sub>±0.74</sub>** | **0.999<sub>±0.001</sub>** | **0.989<sub>±0.007</sub>** | **0.989<sub>±0.007</sub>** |

While the within-task motor-imagery baselines achieve high classification accuracies of 79.20%, 79.91%, and 95.80% on PhysioNet, MULR, and SEMI, respectively. However, SE-RTBNet targets the more challenging ME-to-MI cross-task transfer and mitigates the inherent distribution shift between motor execution and motor imagery signals. Under this cross-task setting, the proposed method maintains competitive over-all decoding performance. Notably, our model outperforms all competing methods across major metrics on the SEMI dataset, attaining an accuracy of 98.94%. These results suggest that the learned feature representations exhibit strong class discriminability and deliver well-rounded performance in terms of classification accuracy and class separation.

Baselines obtain multi-metric advantages under the within-task setting by virtue of task-matched MI training data. SE-RTBNet, nevertheless, still achieves favourable EEG decoding performance under the more demanding ME-to-MI cross-task paradigm.

### C. Subject-Level Accuracy Distribution

To systematically dissect inter-subject variability and comprehensively evaluate the generalization capacity of SE-RTBNet, we visualize the multi-level distribution of subject-wise accuracy across the three datasets, as illustrated in Fig. 2. On the PhysioNet dataset, subject-level classification accuracy ranges from 55% to 97.78%, which objectively reflects the inherent neurophysiological heterogeneity in MI-BCI. Despite pronounced individual differences, most subjects cluster within the interquartile range, and the mean (69.39%±9.41%) closely approximates the median (68.61%). This symmetric, concentrated density profile in the violin plot indicates that SE-RTBNet effectively prevents catastrophic failure for subjects with low-SNR signals and stably extracts discriminative sensorimotor rhythm representations for most subjects.

**Fig. 2.**
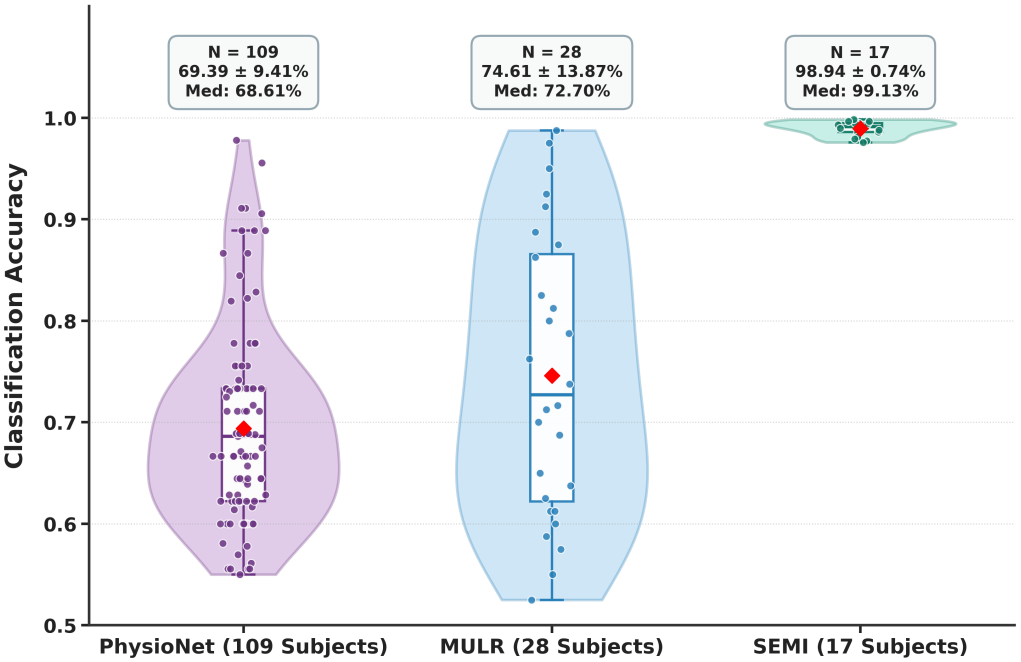
Subject-wise classification accuracy distributions on the PhysioNet, MULR, and SEMI datasets. Violin contours show the kernel density estimation of accuracy, with embedded narrow boxplots depicting interquartile ranges and median lines. Scatter points and red diamonds denote individual subject performance and dataset-level mean accuracy. PhysioNet includes 109 subjects (mean = 69.39%, median = 68.61%), MULR includes 28 subjects (mean = 74.61%, median = 72.70%), SEMI includes 17 subjects (mean = 98.94%, median = 99.13%).

On the MULR dataset, the model achieves a mean accuracy of 74.61% ± 13.87% and a median of 72.70%. The violin plot spans a broad range with an elongated embedded boxplot, indicating notable inter-subject heterogeneity. Nevertheless, the distribution exhibits a prominent high-performance tail, where multiple subjects yield decoding accuracy exceeding 90%.

In sharp contrast, the SEMI dataset yields a smaller variance (0.74) in subject-level performance. Subject accuracies cluster tightly around the global mean of 98.94%, with markedly suppressed inter-individual fluctuations and a highly compressed, flat profile at the upper end of the distribution. Inter-subject performance dispersion is substantially mitigated. Under strictly controlled experimental paradigms with high-SNR recordings, SE-RTBNet reduces prediction bias across subjects and establishes stable, highly discriminative decision boundaries.

### D. Hyperparameter Sensitivity Analysis and Optimal Configuration

To systematically investigate the impacts of key hyper-parameters on classification performance and generalization, a per-subject single-factor control strategy is adopted. For each subject, only the target hyperparameter is adjusted in each experiment while all other architectural and training configurations are fixed, to isolate the individual effect of each parameter. Three representative hyperparameters, including dropout rate (Dropout), shared encoder learning rate (LR_shared_), and latent feature dimensionality (Latent_dim_), are analyzed, corresponding to regularization strength, optimization stability, and feature representation capacity, respectively. All candidate configurations are evaluated via stratified five-fold cross-validation. Validation accuracy serves as the primary screening criterion, and test accuracy is used to assess generalization on unseen data.

Instead of selecting the configuration with the highest test accuracy from a single run, hyperparameter determination follows a multi-criterion selection protocol. First, validation accuracy is used to screen optimal and near-optimal candidates, whose corresponding test accuracy is then examined to evaluate out-of-sample generalization. Among candidates with comparable validation performance, those with higher test accuracy, narrower validation-test accuracy gaps, and lower cross-fold variance are preferred. The final subject-specific optimal configuration is determined by balancing validation performance, test performance, and result robustness, avoiding selection bias from single-run optima.

Based on comprehensive evaluation for each subject, the hyperparameter values highlighted in the pale-yellow regions in Fig. 3 are selected as the final configuration for the corresponding subject in subsequent experiments. This configuration achieves favorable validation performance while maintaining high and stable test accuracy, striking a well-balanced trade-off between model fitting capability and generalization ability.

**Fig. 3.**
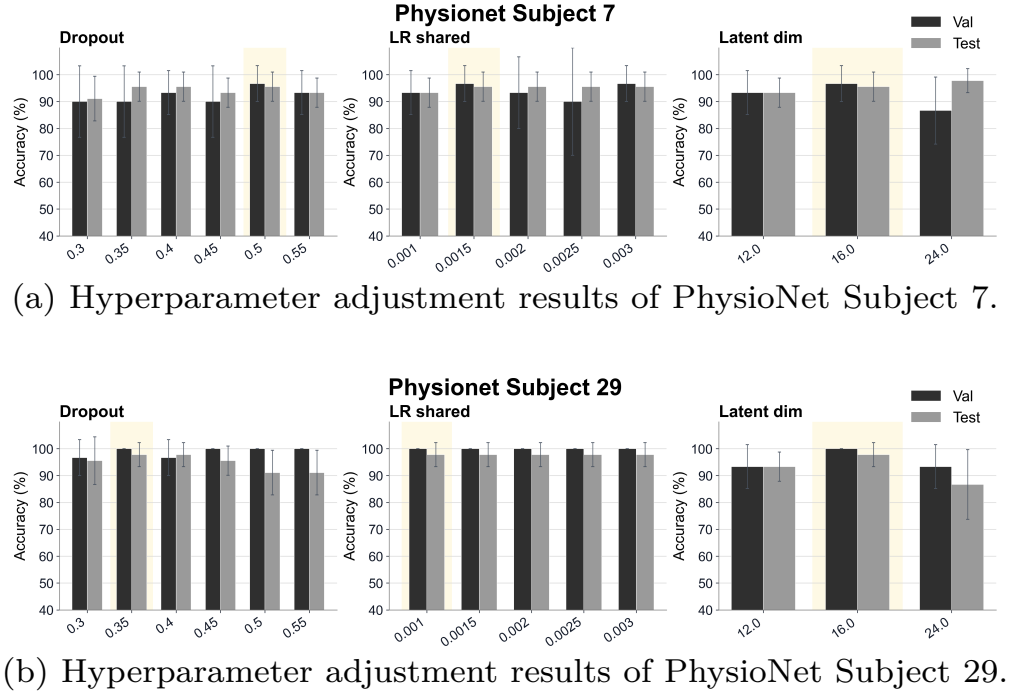
Hyperparameter adjustment results for representative subjects. Dark bars denote validation accuracy, light bars denote test accuracy, and error bars indicate the variation across five folds. The pale-yellow regions mark the final selected hyperparameter values.

## IV. Discussion

### A. Ablation Study

This work conducts ablation studies on the PhysioNet, MULR, and SEMI datasets to validate the functional necessity of each core component embedded in SE-RTBNet. The shared encoder (SharedEnc), Euclidean Alignment (EA), and residual task bridging (Bridge) modules are sequentially removed to construct different model variants. Classification accuracy of these variants is compared with that of the complete SE-RTBNet model. All experimental results are summarized in Table III

**TABLE III.** Ablation study of se-rtbnet on three datasets.

| Model | PhysioNet (%)<br>(*N* = 109) | MULR (%)<br>(*N* = 28) | SEMI (%)<br>(*N* = 17) |
|---|---|---|---|
| w/o SharedEncoder | 57.22<sub>±15.95</sub> | 67.93<sub>±15.82</sub> | 90.26<sub>±6.52</sub> |
| w/o EA | 61.79<sub>±13.82</sub> | 71.00<sub>±16.60</sub> | 97.91<sub>±0.93</sub> |
| w/o Bridge | 60.25<sub>±13.75</sub> | 72.41<sub>±13.85</sub> | 98.61<sub>±0.94</sub> |
| **SE-RTBNet (Ours)** | **69.39<sub>±9.41</sub>** | **74.61<sub>±13.87</sub>** | **98.94<sub>±0.74</sub>** |

The full SE-RTBNet yields the best performance across all three datasets, with classification accuracies of 69.39% ± 9.41%, 74.61% ± 13.87%, and 98.94% ± 0.74%, respectively. Removing SharedEnc leads to the most severe performance drop: accuracy decreases by 12.17 percentage points on PhysioNet, 6.68 percentage points on MULR, and 8.67 percentage points on SEMI. This result demonstrates that the shared encoder serves as the core module for learning general deep representations and enabling cross-domain generalization, which is essential for feature extraction from highly heterogeneous EEG signals.

Removing the EA module leads to accuracy reductions of 7.60, 3.61, and 1.02 percentage points on PhysioNet, MULR, and SEMI, respectively, confirming its effectiveness in mitigating distribution discrepancies. Similarly, removing the Bridge module results in consistent accuracy drops of 9.14, 2.20, and 0.33 percentage points across the three datasets. Although the gains are modest on MULR and SEMI, these consistent improvements highlight the complementary role of residual task bridging beyond shared representation learning and Euclidean alignment. The larger gain on PhysioNet further suggests that the benefits of task-specific compensation vary with cross-task distribution discrepancies. Overall, all three components contribute positively to SE-RTBNet’s performance, supporting their respective roles in the proposed framework.

### B. EEG Channel Saliency Interpretation

To verify whether SE-RTBNet captures neurophysiologically meaningful patterns, we adopt EEG saliency topomaps to visualize channel importance. Unlike raw amplitude or ERD/ERS maps, these saliency maps (Fig. 4) quantify scalp-region contributions to MI predictions and serve as qualitative importance indicators instead of direct cortical activation measurements. As discriminative EEG signatures for ME/MI mainly lie over sensorimotor regions (C3, Cz, C4), our maps assign greater weights to spatially contiguous centro-lateral and sensorimotor areas, matching known rhythmic modulations during hand imagery [12]. Variations across samples are anticipated owing to inter-subject, inter-trial and cross-dataset EEG variability.

**Fig. 4.**
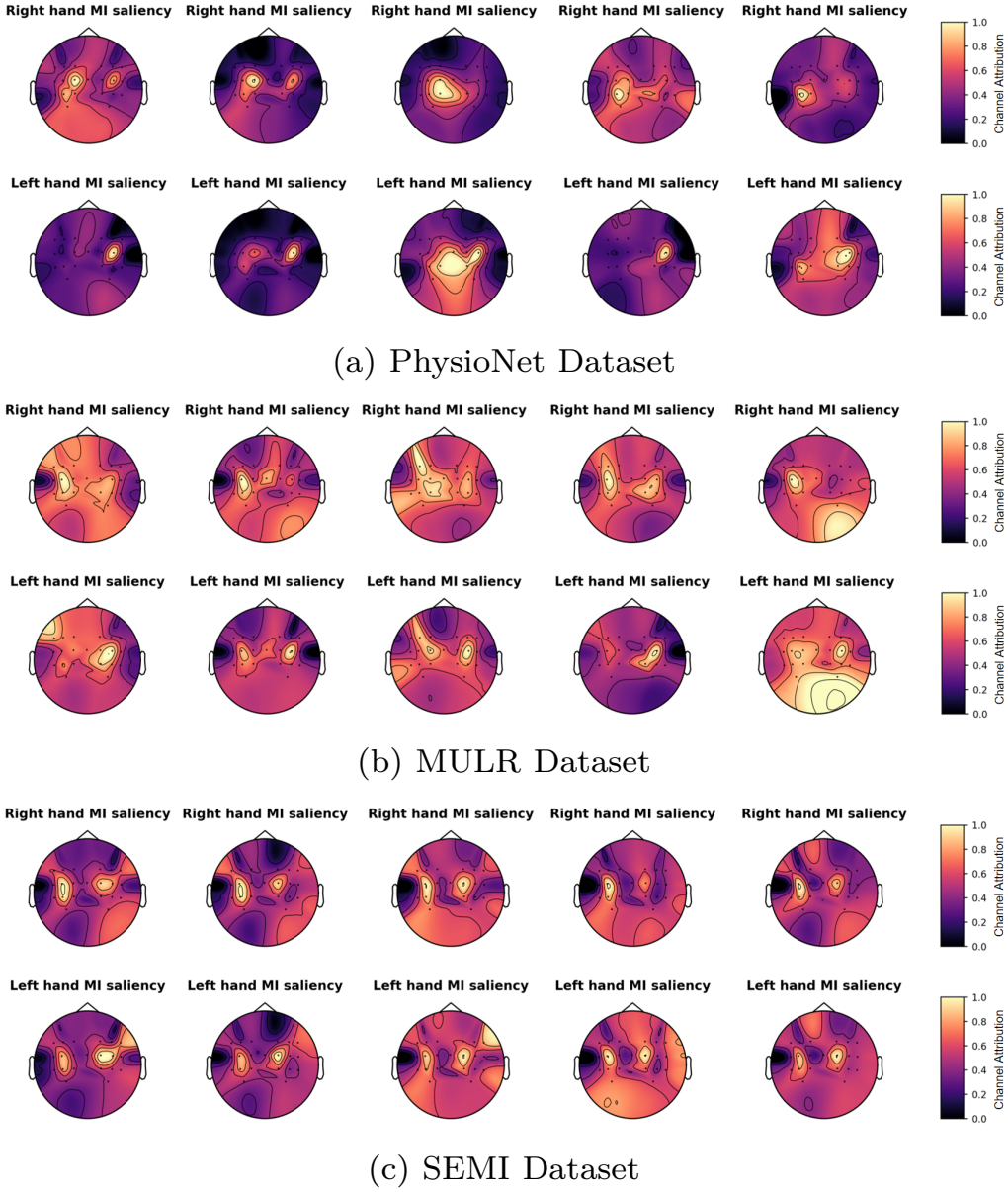
Representative MI saliency topographic maps of SE-RTBNet across three datasets: (a) PhysioNet, (b) MULR Dataset, and (c) SEMI Dataset. For each dataset, the scalp maps present the spatial importance distributions for right-hand and left-hand motor imagery, with warmer colors indicating higher model-assigned feature importance on the normalized scale. The maps provide qualitative evidence that SE-RTBNet relies on structured spatial information rather than arbitrary channel responses.

These interpretability findings supplement quantitative classification results. The structured, partly asymmetric saliency patterns indicate the model learns spatially meaningful motor-related EEG features rather than random noise. This topo-graphic evidence also clarifies the advantage of the shared encoder: projecting ME and MI samples into a common feature space boosts classification accuracy and produces features with higher neurophysiological plausibility.

### C. Latent Feature Distribution and Cross-Task Alignment

To illustrate how SE-RTBNet mitigates ME-to-MI domain shift, Fig 5 shows 2D t-SNE projections across three model stages: the shared encoder space (a), residual task bridging (b), and final prototypical inference space (c).

**Fig. 5.**
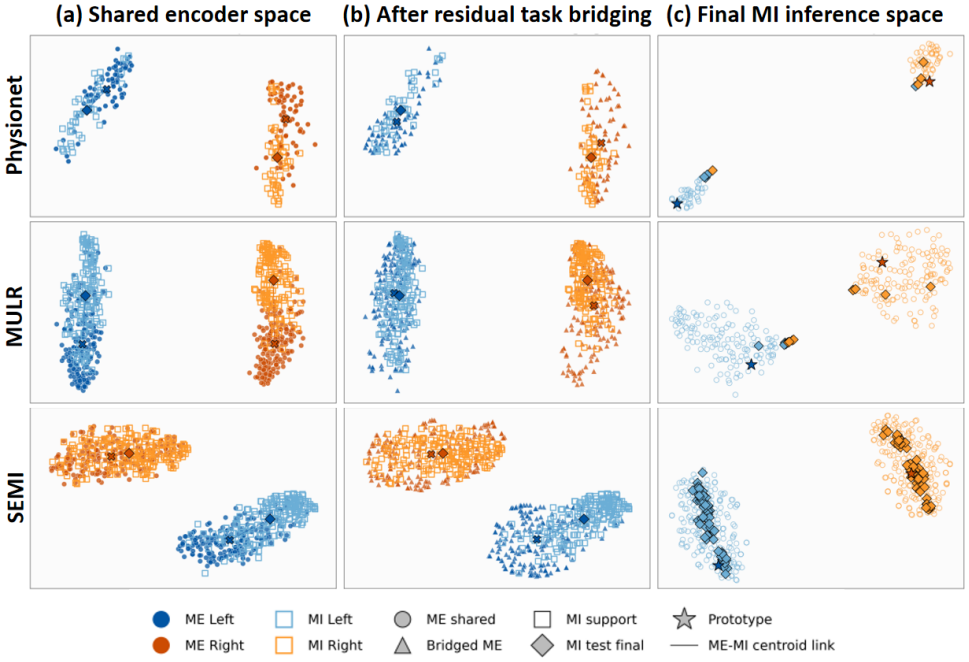
Latent feature space evolution across three model stages evaluated on the PhysioNet (top row), MULR (middle row), and SEMI (bottom row) datasets: (a) Initial space from the shared encoder, (b) Bridged feature space via the residual task-bridging module, and (c) Calibrated target MI inference space.

In the shared encoder space (a), ME and MI features exhibit distinguishable class structures but retain notable distributional differences. After residual task bridging (b), bridged ME features move closer to their corresponding MI support distributions, indicating improved alignment. This effect is more evident on PhysioNet and MULR, while SEMI retains some residual discrepancies, reflecting dataset-dependent alignment difficulty. Nevertheless, in the final MI inference space (c), adaptive prototype calibration and test-time augmentation produce distinguishable class-wise distributions with clear inter-class separation across all datasets. These results demonstrate that SE-RTBNet mitigates cross-task discrepancies while preserving discriminative structures for MI inference.

### D. Spatio-Temporal Attribution via Integrated Gradients

We calculate spatiotemporal attribution maps on the test set via the Integrated Gradients algorithm to verify whether discriminative representations extracted by SE-RTBNet conform to canonical sensorimotor rhythm neurophysiological mechanisms, as illustrated in Fig 6.

**Fig. 6.**
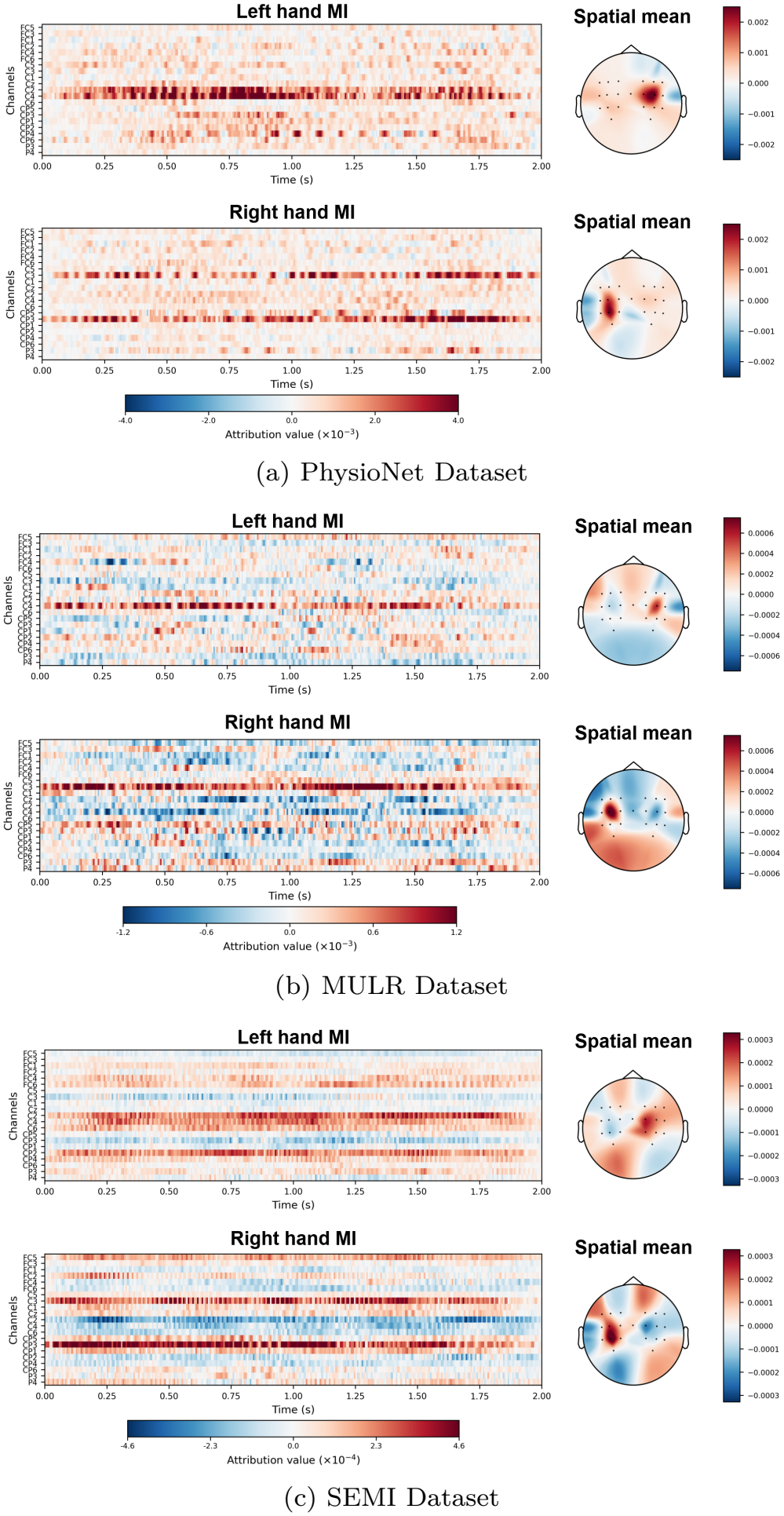
Spatiotemporal attribution visualization of SE-RTBNet across three benchmark datasets: (a) PhysioNet, (b) MULR, and (c) SEMI. For each dataset, the temporal heatmaps illustrate feature attribution dynamics across motor channels for left- and right-hand motor imagery, while the corresponding scalp topographies display the spatial mean attribution distributions. Positive attribution values represent features that contribute positively to target-class decisions, whereas negative values reflect suppressive or countervailing evidence.

Spatially, the scalp topographies exhibit pronounced contralateral hemispheric dominance, which is highly consistent with classic ERD/ERS patterns [40]. During left-hand motor imagery (Left hand MI), prominent positive attributions are densely concentrated over the contralateral right sensorimotor cortex, centered primarily on channel C4 and adjacent centroparietal electrodes. Conversely, during right-hand motor imagery (Right hand MI), the focal activation shifts distinctly toward the contralateral left motor area. This sharp cross-hemispheric lateralization demonstrates that the decision-making process of the proposed model relies on genuine contralateral cortical neurodynamics rather than non-specific background artifacts.

Temporally, the feature attributions reveal dynamic evolution across the 2.0-second time window. The discriminative energy is not uniformly distributed. Instead, it exhibits rhythmic bursts predominantly between 0.75 s and 1.50 s, which corresponds to the peak phase of mental motor preparation and sustained imagery execution.

Furthermore, the physiological prior loss incorporated during training regularizes the spatial attention weights, ensuring that the residual task-bridging module aligns the ME and MI domains while strictly preserving the intrinsic sensorimotor topology. In summary, the attribution analysis substantiates that the superior performance of SE-RTBNet is grounded in sound neuroanatomical principles and highly interpretable representation learning.

### E. Model Label Sensitivity and Transfer Efficiency

The experimental results under different ratios of labeled MI samples further demonstrate the label efficiency of SE-RTBNet in ME-to-MI transfer learning, as shown in Fig. 7. Across the three datasets, the classification accuracy of the model rises steadily as the proportion of labeled MI samples increases from 20% to 80%. The significance markers (^***^) indicate that expanding MI annotations yields statistically significant performance improvements. Even with only 20% MI labels, the model achieves acceptable decoding accuracy, verifying that discriminative knowledge learned from ME data can compensate for the scarcity of MI annotations.

**Fig. 7.**
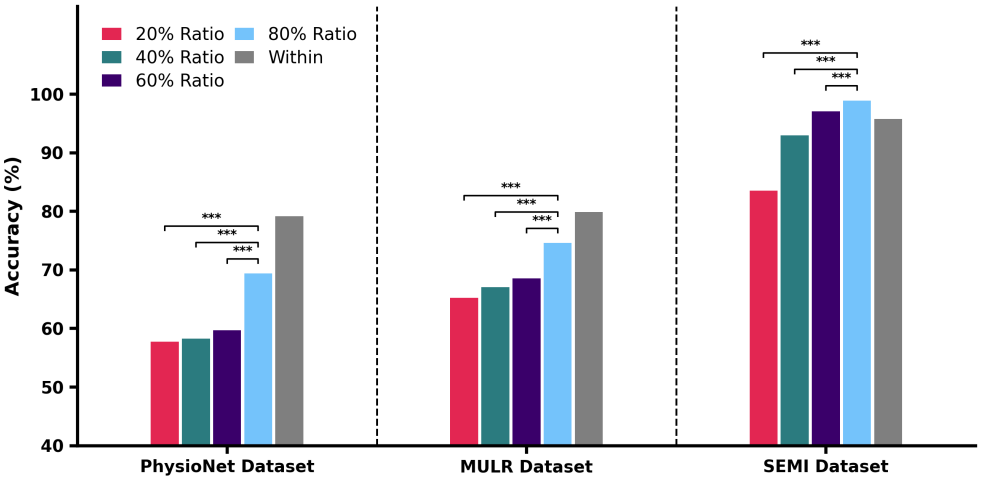
Classification accuracy of SE-RTBNet under different ratios of labeled MI samples across three datasets. “Within” denotes the within-task MI-train-MI-test results of EEGConformer as an external reference. Asterisks ^***^ indicate statistically significant differences (***p <* 0.001**).

A consistent performance gap persists between the transfer accuracy under the 80% label setting and the within-task “Within” baseline on the PhysioNet and MULR datasets. This finding reveals that although ME features provide valuable auxiliary cues for MI classification, the inherent distribution shift of neural representations between motor execution and motor imagery cannot be fully resolved merely by increasing target-domain labels. In sharp contrast, SE-RTBNet exhibits outstanding capability for representation alignment and performance saturation on the high-quality SEMI dataset, where the transfer performance with 80% labeled samples surpasses the within-task baseline. This comparative result strongly demonstrates that the proposed cross-task transfer architecture can adequately exploit shared sensorimotor rhythm representations to match or even approach the performance upper bound of conventional within-task MI decoding.

### F. Limitations and Future Works

Although SE-RTBNet shows promising performance for ME-to-MI transfer learning, several limitations remain. First, the improvement varies across datasets. The method achieves promising results on the PhysioNet and MULR datasets quality, and its classification accuracy on the SEMI dataset exceeds that of the within-task MI baseline. This suggests that dataset characteristics, such as subject variability, task consistency, and signal quality, can strongly influence the observed effectiveness of transfer learning methods. Future work should evaluate the model on more diverse EEG datasets to better characterize its generalization ability.

Second, the current study focuses on binary left-hand versus right-hand classification. While this setting is widely used in motor imagery research, practical BCI systems often require more complex control paradigms. Extending SE-RTBNet to multi-class and more realistic online BCI scenarios would be an important direction for future research.

Finally, the current framework relies on offline preprocessing and evaluation. While this setup facilitates systematic bench-marking, real-world BCI applications demand models robust to session-to-session variability, nonstationary EEG signals, and limited calibration data. Future studies may explore online adaptation strategies, lightweight calibration mechanisms, or subject-specific updating schemes to improve deployment feasibility.

## V. Conclusions

This paper presents SE-RTBNet, a Shared-Encoder Residual Task-Bridge Network for ME-to-MI cross-task transfer learning in EEG-based left/right hand motor imagery decoding. SE-RTBNet learns shared sensorimotor representations from multi-band EEG signals and introduces a residual task-bridging mechanism to project ME features into the MI latent space, preserving discriminative source-task information while compensating for the representation gap between actual movement and imagery. Experiments on three datasets verify the framework’s effectiveness. Under the ME-trained/MI-tested protocol, SE-RTBNet achieves 69.39%, 74.61%, and 98.94% accuracy on PhysioNet, MULR, and SEMI, respectively, outperforming state-of-the-art methods. SE-RTBNet holds the potential to significantly shorten the lengthy and fatiguing calibration training cycles for patients with movement disorders, offering significant clinical value and broad application prospects for practical translation in areas such as neurorehabilitation, the restoration of active motor function, and assistive exoskeleton control.

